# The osteoarthritis associated sphingolipid sphingomyelin 34:1 causes inflammatory pain in mice

**DOI:** 10.1101/2024.11.19.621807

**Authors:** Rebecca H. Rickman, Luke A. Pattison, Lanhui Qiu, Maya Dannawi, Fraser Welsh, Ewan St. John Smith

## Abstract

**Objective:** Osteoarthritis (OA) has a multifactorial pathogenesis, pain being the main symptom driving clinical decision making. Dysregulation of multiple mediators occurs in OA, the roles of many remaining to be identified. In dogs and humans with OA, synovial fluid lipidome dysregulation occurs, some findings being replicated in the plasma lipidome in a mouse OA model. One upregulated lipid is the sphingomyelin N-palmitoyl-D-erythro-sphingosylphosphorylcholine (d18:1/16:0), referred to here as SM. This study aimed to determine if SM causes joint pain and neuronal hyperexcitability in mice.

**Design:** The effects of SM or a structurally related ceramide (CM) on mouse sensory neuron excitability were measured using patch-clamp electrophysiology, as well as the ability of intraarticular SM and CM to induce inflammatory pain in mice.

**Results:** Incubation of sensory neurons with 1 µM SM decreased rheobase, compared to incubation with vehicle (*p-adj* = 0.0000146, 95% confidence interval (CI): 50.20, 76.73) or CM (*p-adj* = 0.138, CI: 103.45, 171.55). Similarly, SM induced mechanical hypersensitivity in mice compared to mice receiving vehicle (*p-adj =* 0.000003, 95% confidence interval (CI): 166.82, 251.63) or CM (*p-adj =* 0.055, 95% CI: 218.28, 268.12), which was coupled with a significant decrease in rheobase of knee-innervating neurons isolated from SM-injected mice compared to those receiving vehicle *(p-adj =* 0.0138, CI: 50.19, 76.73) or CM (*p-adj* = 1.0, CI: 103.45, 171.55).

**Conclusions:** The results generated demonstrate that a dysregulated lipidome can contribute to inflammatory OA pain, further work being necessary to determine the mechanism by which SM exerts its activity.

## Introduction

Osteoarthritis (OA) is a chronic degenerative condition of synovial joints associated with joint stiffness, swelling, and pain, which drives clinical decision making^1^. OA pathogenesis and pain mechanisms are multifactorial, but joint inflammation underpins pain sensitisation in OA^2^. Knee-innervating sensory neurons interact with multiple cell types and mediators that can activate and/or sensitise neurons^3^. Synovial fluid from OA individuals, but not individuals without joint disease, sensitises murine knee-innervating neurons^4^. Identifying those mediators modulating sensory neuron excitability could identify new therapeutic opportunities that are much needed considering the inadequacy of current OA pain management strategies. Among the plethora of pain-inducing mediators, analysis of the synovial fluid lipidome of humans and dogs with OA has identified dysregulation of numerous lipids^5,6^. Furthermore, in a mouse model of OA, analysis of plasma lipid levels identified differential expression of 24 lipids, levels of the sphingomyelin N-palmitoyl-D-erythro-sphingosylphosphorylcholine (d18:1/16:0), also known as SM(d34:1), correlating with cartilage damage and trending towards correlations with changes in pain behaviour^7^. This sphingomyelin, SM(d34:1), was also upregulated in the synovial fluid of OA dogs and humans^5,6^. The structurally similar ceramide, N-palmitoyl-D-erythro-sphingosine (d18:1/16:0), was also upregulated in human OA synovial fluid^5^, but not detected in canine samples^6^, and was not upregulated in a mouse OA model^7^. Lipids are well-characterised for their ability to modulate neuronal excitability, for example, exudates from individuals with acute knee-joint effusion have high levels of arachidonic acid and lysophosphatidylcholine, which both activate and sensitise acid-sensing ion channel 3 (ASIC3)^8,9^. Considering the cross-species OA-associated upregulation of SM(d34:1), we sought to determine if SM(d34:1) contributes to OA joint pain in mice.

## Methods

Further methods and data analysis is available in the online supplemental file.

### Animals

C5BL/6J mice (Envigo) were housed in groups of up to 5/cage with free access to food and water. Holding rooms were maintained at 21 °C and operated a 12-hour light/dark cycle. Animal work was regulated in accordance with the United Kingdom Animal (Scientific Procedures) Act 1986 Amendment Regulations 2012. For behavioural studies 44 female mice aged 10-12 weeks were used, 8 of which were used postmortem for electrophysiology analysis of knee-innervating neurons. For in vitro electrophysiology, 7 males and 5 females (10-15 weeks) used.

### DRG neuron culture

As described previously^4,10,11^, lumbar DRG (L2-L5) were isolated from humanely killed mice, incubated in collagenase (3mg, 15 min; Merck) and then trypsin (3mg, 30 min; Merck). Following dissociation with mechanical trituration, neurons were plated on poly-D-lysine and laminin glass-bottomed dishes (MatTek).

### Electrophysiology

DRG neurons were bathed in extracellular solution (ECS) (in mM): NaCl (140), KCl (4), CaCl_2_ (2), MgCl_2_ (1), glucose (4), HEPES (10), adjusted to pH 7.4 with NaOH, and osmolality adjusted to 280-295 mOsm with sucrose. Intracellular solution (in mM) contained KCl (110), NaCl (10), MgCl_2_ (1), EGTA (1), and HEPES (10), adjusted to pH 7.4 with KOH and an osmolality of 300-310 mOsm. Data were acquired using a Multiclamp 700A amplifier and Digidata 1440A digitizer (Molecular Devices). Patch pipettes (3-6 MΩ) were pulled using a P-97 Flaming/Brown puller (Sutter Instruments) from borosilicate glass capillaries (Hilgenberg). Capacitance of cells from different groups was similar (Supplementary Tables 1-3). Current was injected (-100 pA to 2000 pA, 80 ms, 21 steps) to generate action potentials, data sampled at 20kHz and filtered at 5kHz. N-palmitoyl-D-erythro-sphingosylphosphorylcholine, SM(d34:1) and N-palmitoyl-D-erythro-sphingosine, CM(d34:1) were purchased from Avanti Polar Lipids (860584 and 860516). Chloroform and methanol (Sigma, 2883069 and 32213) were mixed at an 8:2 ratio, lipids dissolved in a 0.1% chloroform:methanol solution (also used as a vehicle control). For overnight incubation experiments, neurons were cultured with lipids at a final concentration of either 1 µM or 100 µM or 0.001% / 0.1% chloroform:methanol for 18-24-hours before recordings. For recordings from Fast Blue labelled neurons (see below), neuronal isolation occurred 6-hours after lipid/vehicle injection, recordings taking place 10-17-hours after plating. The experimenter was blind to experimental conditions.

### Intra-articular injections

Labelling of knee-innervating sensory neurons was achieved by intra-articular injection of Fast Blue (1.5 µl, 2% (w/v) in saline; Polysciences) under anaesthesia (ketamine, 100 mg/kg and xylazine, 10 mg/kg, i.p.) 1-week before injection of lipids/vehicle. Intra-articular injections of 5 µl 1 µM SM(d34:1), CM(d34:1), or 0.1% chloroform:methanol (8:2) prepared in saline were made under inhalation anaesthesia (isoflurane: 4% induction, 2% maintenance) unilaterally, to a randomly determined knee (using customised scripts written in R), mice kept in a recovery chamber following injections. Knee widths were measured with digital callipers before, 6- and 24-hours post-injections.

### Behavioural Analyses

Behavioural experiments were performed in the presence of a female and male experimenter. Mice were acclimatised to the testing room for at least 30-minutes before behaviours were tested. When assessing multiple behaviours, order of testing followed: digging, RotaRod and pressure application measurement.

### Behaviour – Pressure Application Measurement

Knee joint mechanical sensitivity was assessed using a pressure application measurement device (Ugo Basile). Animals were scruffed by 1 experimenter and presented to a second experimenter, who was blind to the animal’s identity. The second experimenter applied increasing force to the ipsilateral knee joint medially with the force transducer, withdrawal threshold being recorded when an animal withdrew the limb or when reaching 450g of force. Each animal was tested twice per time point, with a short break between tests, withdrawal threshold reported as an average of the 2 measurements.

### Behaviour – Digging

Digging behaviour was assessed as a readout of how pain affects an ethologically relevant mouse behaviour^12^. Testing involved placing individual animals in cages containing ∼4cm of tightly packed fine-grain aspen midi 8.20 wood chip bedding (LBS Technology). Mice were allowed 3-minutes to explore without interference. Mice were first placed in test cages the day before baseline behaviour measurement, to gain familiarity with the testing environment. Test sessions were video-recorded, after the 3-minutes the number of visible burrows remaining was recorded. Digging behaviour was further analysed at the conclusion of studies, two experimenters recording the time spent digging in videos after blinding and an average of the time reported as the time spent digging.

### Behaviour – Rotarod

A RotaRod (Ugo Basile) was used to assess the motor coordination of mice. Animals acclimatised for 1-minute to the apparatus on a slow setting (7 rpm) before initiating an accelerating program (7-40 rpm, 5-minutes). Mice were first placed on the Rotarod the day before baseline behaviour measurement, to gain apparatus familiarity. Test sessions were video-recorded, and mice were removed from the RotaRod if they fell, after 2 consecutive passive rotations, or after 6 minutes of the accelerating program. Video analysis was performed at the conclusion of studies, after blinding, latency to passive rotation or fall being scored by a single experimenter.

### Data Analysis

Number of animals used per experiment is detailed in the corresponding figure legend; no inclusion/exclusion criteria used. Data are presented as mean ± standard error of the mean; raw data supporting this study are available: https://doi.org/10.17863/CAM.113300. Appropriate analyses were selected according to the number of factors being compared and whether data met the assumptions for parametric analyses – statistical tests are detailed in figure legends, analyses performed in R, and *p* values less than 0.05 were considered significant.

## Results

Pain is initiated via sensory neuron activation. Therefore, to simulate the OA joint, mouse DRG neurons were cultured overnight in the sphingomyelin SM(d34:1), the structurally similar ceramide CM(d34:1), or chloroform:methanol as a vehicle control; hereafter referred to as SM, CM and Veh, Fig. 1A. In humans, the approximate ranges reported are 150 nM – 1 µM and 10 µM – 100 µM for CM and SM respectively^5^. Compared to Veh, 1 µM SM, but not 1 µM CM, depolarised the resting membrane potential and decreased rheobase, i.e. neuronal sensitisation, an effect consistent in neurons isolated from male and female mice (Fig. 1B and C); there was limited impact on other action potential parameters (Table S1). At 100 µM, both SM and CM induced sensitisation (Fig. S1 and Table S2). However, because 100 µM is beyond the range observed in vivo for CM^5^ we decided to examine the ability of CM/SM to induce inflammatory joint pain in mice using 1 µM. Due to the increased prevalence of OA in females^1^ and the lack of sex effect observed with SM in vitro, we only conducted in vivo experiments in females. Each substance was intra-articularly injected into one knee joint, swelling and behavioural measurements made 6- and 24-hours post-injection (Fig. 1D). Whereas no change from baseline knee width occurred in Veh or CM groups, SM significantly increased knee width, being most pronounced 6-hours post-injection, but persisting 24-hours post-injection (Fig. 1E). Concomitant with knee inflammation, SM injection decreased mechanical withdrawal threshold, an evoked pain measurement that provides readout of primary hyperalgesia (Fig. 1F). In addition, SM decreased the amount of time mice spent digging and number of burrows dug, a readout of how ongoing pain affects a natural behaviour of mice (Fig. 1G and H). However, SM did not impact time spent on the rotarod, i.e. SM did not negatively impact motor function (Fig. 1I). As might be expected from the lack of inflammation, CM produced no significant change in animal behaviour, but, unexpectedly, Veh injection did cause a mild, yet significant, decrease in mechanical withdrawal threshold.

**Figure 1.**
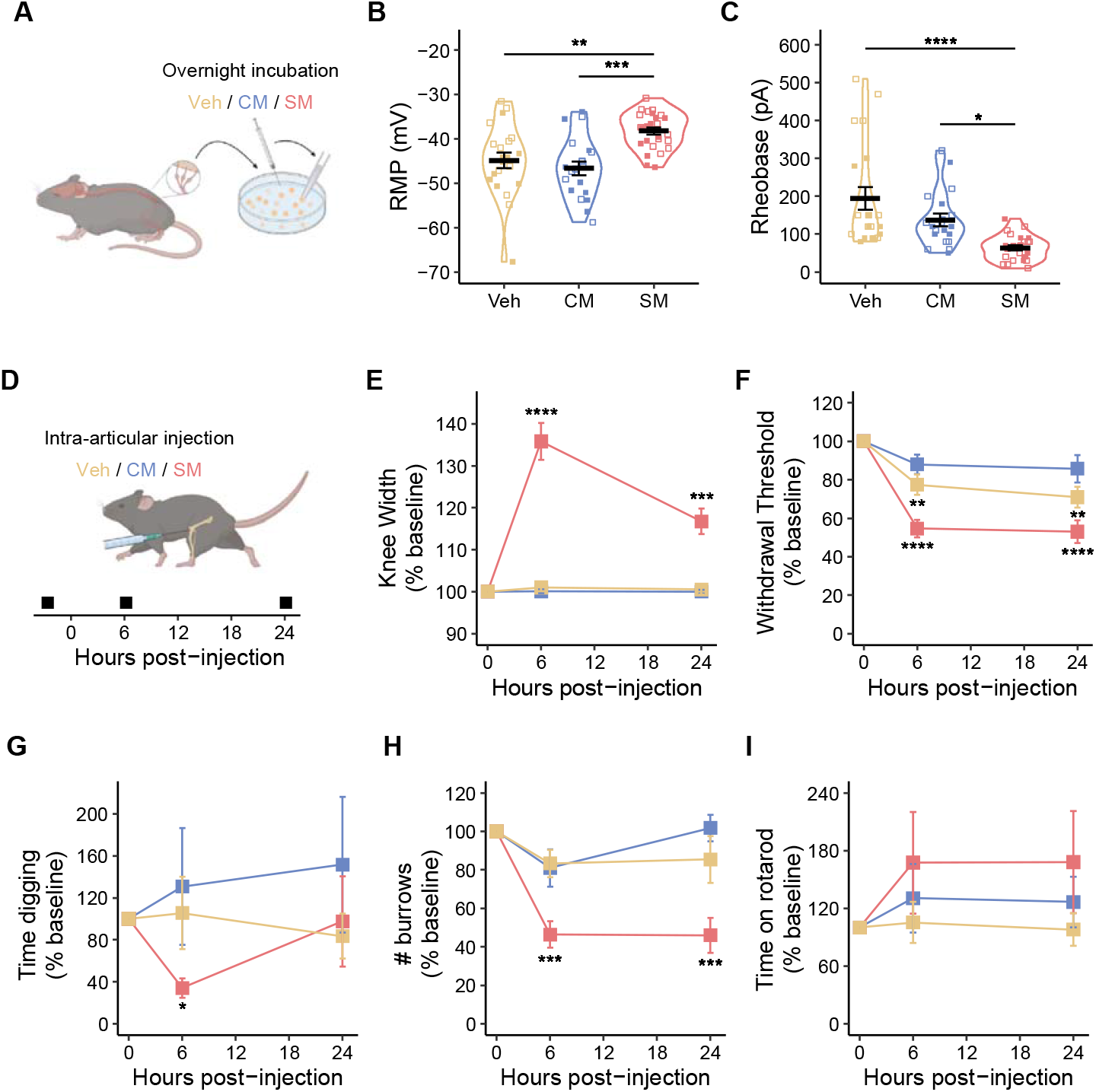
SM induces neuronal hyperexcitability and inflammatory pain *in vivo*. **(A)** Näive sensory neurons were incubated with either vehicle (Veh), 1 µM (CM) or 1 µM (SM) overnight before the effects on **(B)** resting membrane potential (RMP) and **(C)** rheobase were assessed by whole-cell patch clamp electrophysiology. **(D)** After capturing baseline knee widths and behaviours, mice received a unilateral intra-articular injection of 5 µl Veh, 1 µM CM or 1 µM SM, **(E)** knee-width, **(F)** mechanical sensitivity of the inject knee, **(G)** time spent digging, **(H)** number of burrows produced and **(I)** time on the Rotarod were re-assessed 6- and 24-hours post injections. * *p-adj* < 0.05, ** *p-adj* < 0.01, *** *p-adj* < 0.001, **** *p-adj* < 0.0001: **(B-C)** One-way ANOVA with Bonferroni-correct post hoc, n = 21 (Veh), 20 (CM), 26 (SM) cells, N = 4 (2 male, open symbols, and 2 female, closed symbols); **(E-I)** Repeated-measures ANOVA with Bonferroni-correct post hoc, annotated differences are the effect of testing point for each injection substance, N = 12 per injection substance.

Considering that 1 µM SM, but not CM, induced inflammatory joint pain (Fig. 1E-I), but that overnight incubation of DRG neurons with only high concentration (100 µM) of SM or CM caused neuronal sensitisation (Fig. S1), a possible explanation is that 1 µM SM can initiate an in vivo sensitisation signalling cascade that is not activated by 1 µM CM. To determine if this was the case, knee-innervating neurons in mice were labelled with Fast Blue before 1-week later administering an intra-articular injection of SM/CM/Veh and then 6-hours later measuring knee width and digging behaviour before isolating neurons for electrophysiology analysis (Fig. 2A). The experimental endpoint of 6-hours was chosen because this was the timepoint at which greatest inflammation had previously been observed (Fig. 1E). As expected, in this smaller cohort of mice, again, only SM induced knee inflammation (Fig. 2B) and decreased digging behaviour (Fig. 2C). Electrophysiology recordings of knee-innervating neurons demonstrated that SM induced a significant depolarisation of the resting membrane potential and a significant decrease in rheobase compared to knee-innervating neurons isolated from CM or Veh injected mice (Fig. 2D and E); other action potential parameters not being significantly affected (Supplementary Table 3).

**Figure 2.**
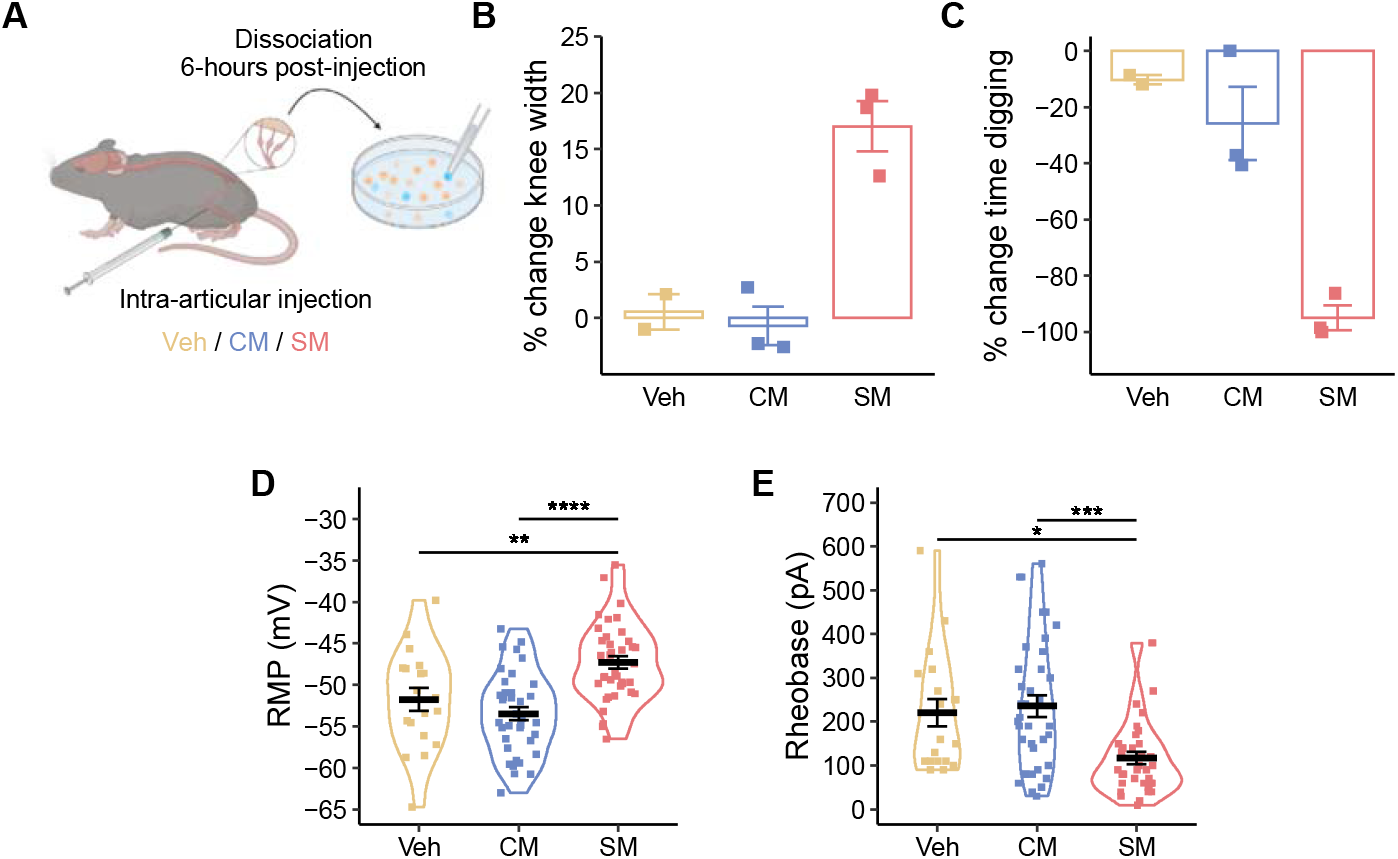
Knee-innervating neurons are hyperexcitable following intra-articular injection of SM. **(A)** Knee-innervating neurons of mice were labelled with the retrograde tracer Fast Blue, a week later mice received a unilateral injection of either 5 µl Veh, 1 µM CM or 1 µM SM. The effect of injections on **(B)** knee width and **(C)** time spent digging was assessed 6-hours post-injection, after which mice were euthanised and sensory neurons collected and cultured for electrophysiology experiments. The effect of intra-articular injections on **(D)** resting membrane potential (RMP) and **(E)** rheobase was assessed focussing on Fast Blue-positive cells. * *p-adj* < 0.05, ** *p-adj* < 0.01, *** *p-adj* < 0.001, **** *p-adj* < 0.0001: **(B-C)** N = 2 (Veh), 3 (CM and SM); (**D-E**) One-way ANOVA with Bonferroni-correct post hoc, N = 8 (all female) n = 19 (Veh), 37 (CM), 38 (SM) neurons.

## Discussion

In OA, synovial fluid contents are altered due to cartilage breakdown and secretion of mediators by numerous cell types^1,3^. In recent years, there has been much focus on fibroblast-like synoviocytes, cells whose phenotype is altered in painful OA^13^ and cells that can respond to OA-relevant stimuli to release inflammatory mediators, which in turn sensitise knee-innervating neurons^10,11^. It has been demonstrated that the synovial fluid and plasma lipidome is dysregulated in OA^5–7^ and this study investigated the ability of a sphingomyelin, which is upregulated across species in OA^5–7^, to cause inflammatory knee pain. Overnight incubation of DRG neurons with 1 µM SM but not CM, sensitised neuronal function, whereas 100 µM of either lipid caused sensitisation. In vivo experiments were conducted using 1 µM SM/CM due this being at the upper end of [CM] reached in vivo in the synovial fluid from humans with late stage OA^5^. In line with in vitro results, only SM induced knee inflammation and pain behaviour. CM and SM are both sphingolipids, replacement of a hydroxy group with a phosphorylcholine resulting in a sphingomyelin rather than a ceramide, thus, both lipids share heavily saturated acyl chains. Lipid modulation of ion channels is a well-described phenomenon, involving either direct lipid-protein interactions or indirect modification of membrane properties^14^. One feature of plasma membranes and ion channel modulation is lipid rafts, small, sphingolipid enriched membrane domains, disruption of which has been proposed as a potential method of providing analgesia^15^. Therefore, one explanation for both 100 µM SM and CM causing neuronal sensitisation following overnight incubation is that they modulate the plasma membrane lipid raft composition in a similar manner, which alters ion channel function to produce a similar degree of sensitisation. The fact that 1 µM SM, but not 1 µM CM, induces neuronal sensitisation in vitro could be due to a more potent impact on plasma membrane lipid rafts, or, additionally and/or alternatively, SM might initiate signalling events that CM cannot.

The fact that only SM caused inflammatory pain in mice and that knee-innervating neurons from SM, but not CM injected animals, were hyperexcitable indicates that SM likely coordinates signalling events in vivo that CM cannot. Considering that inflammation is observed alongside neuronal hyperexcitability, a plausible explanation is that SM stimulates a non-neuronal cell type to drive an inflammatory response, which includes sensitisation of knee-innervating neurons by pro-inflammatory mediators. Further work is required to determine the mechanisms involved in SM-mediated inflammation, potentially involving activation of an as yet unknown receptor, or perturbation of lipid-mediated signalling in an SM-dependent manner, as well as examining the persistence of SM-induced inflammatory pain. We cannot rule out that higher concentrations of CM when delivered in vivo would also cause inflammatory pain, but we did not pursue this due to 1 µM being reported as the upper concentration of CM reached in the synovial fluid from humans with late stage OA^5^.

In summary, this study demonstrates that the elevated levels of a specific sphingomyelin (sphingomyelin N-palmitoyl-D-erythro-sphingosylphosphorylcholine (d18:1/16:0)) observed in OA likely contribute to OA pain. This is demonstrated by administration of SM, but not the structurally related ceramide (N-palmitoyl-D-erythro-sphingosine (d18:1/16:0)) inducing swelling, pain-related behaviours and knee-innervating neuron hyperexcitability. This study thus highlights the necessity of understanding the biology of those molecules upregulated in OA synovial fluid with further work being required to determine the precise mechanism of SM-mediated inflammatory pain and thus determine the potential druggability of this pathway.

## Acknowledgments

We thank Combined Animal Facility technical staff for their assistance and Profs. Nicolas Cenac and Thierry Levade for discussing preliminary results.

## Author contributions

RHR, LAP, LQ and MD contributed equally, author order reflects timeline of project participation. Conception and design (RHR, LAP, LQ, MD, FW and EStJS), Collection and assembly of data, (RHR, LAP, LQ, and MD), Analysis and interpretation of the data (RHR, LAP, LQ, MD, FW and EStJS), Drafting and final approval of the article (RHR, LAP, LQ, MD, FW and EStJS), and Obtaining of funding (FW and EStJS).

## Supplementary Methods, Figure and Tables

### Animals

The table below provides an overview of the animals used in each electrophysiology experiment and the number of neurons recorded from per mouse.

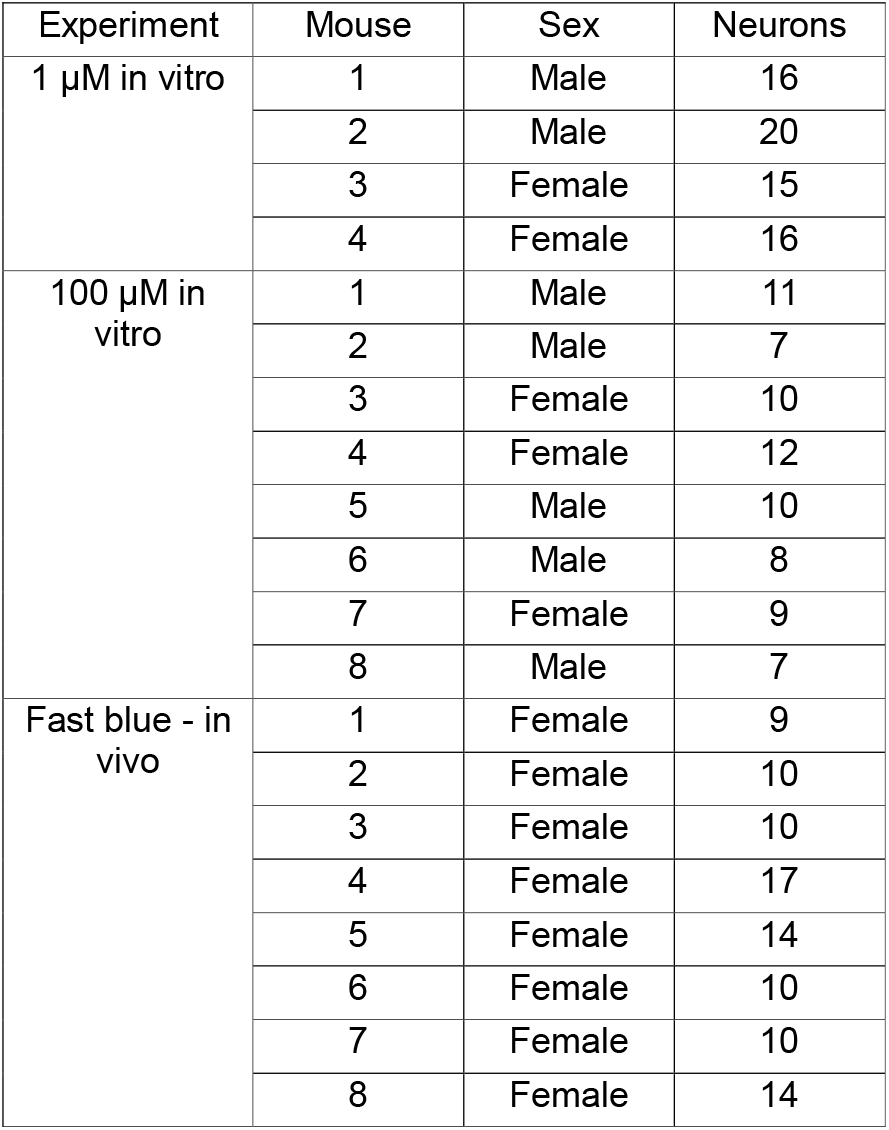

### Study Design and Statistical Analysis

For in vitro electrophysiology experiments involving overnight incubation with different substances, the independent biological unit for comparison was the neuron. Neurons isolated from each mouse were plated in 6 dishes and vehicle, CM or SM applied randomly to 2 dishes each, the experimenter being blinded as to whether neurons were incubated overnight with vehicle, CM or SM. Multiple outcome variables were assessed: resting membrane potential, rheobase, capacitance, and action potential amplitude, half-peak duration, afterhyperpolarisation amplitude and duration. All data were scrutinised to verify that they met the assumptions of parametric analyses. Normality was assessed using the Shapiro-Wilk test. One-way ANOVA with Bonferroni-correct post hoc analysis was conducted to measure differences between groups using R, *p* values less than 0.05 being considered significant. The same method was conducted for analysis of fast blue labelled, knee-innervating neurons, the experimenter being blinded as to whether neurons were isolated from vehicle, CM or SM treated mice.

For behavioural experiments, the independent biological unit for comparison was the mouse, a randomisation script in R being used to assign mice to different groups (vehicle, CM and SM). The experimenters were blinded to whether animals were injected with vehicle, CM or SM. Multiple outcome variables were measured: knee width, mechanical withdrawal threshold, time spent digging, number of burrows dug, and time spent on the rotarod. Data were deemed normally distributed, through Shapiro-Wilk tests and thus repeated-measures ANOVA with Bonferroni-correct post hoc test was used to measure differences between treatment groups over time using R, *p* values less than 0.05 being considered significant.

## Supplementary Figure

**Figure S1.**
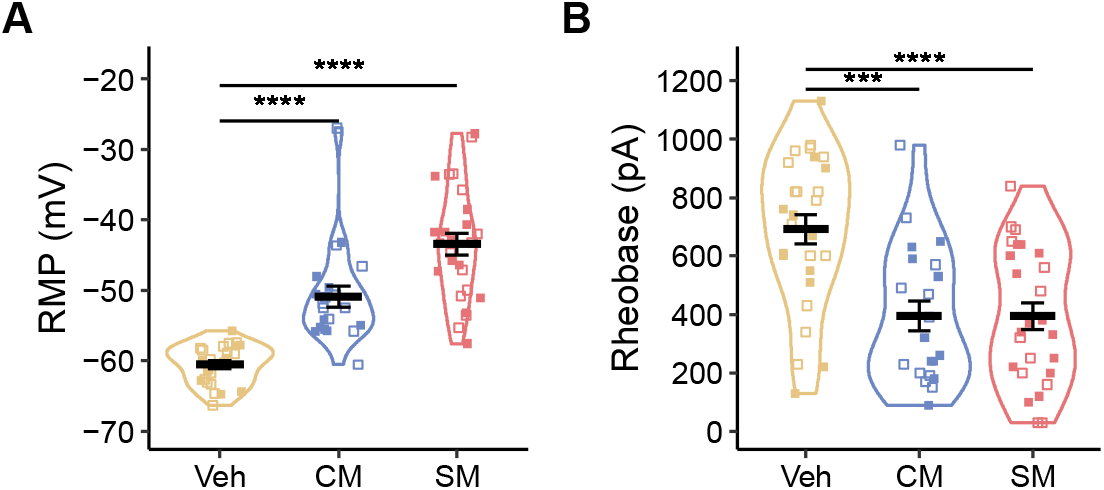
100 µM SM and CM induce neuronal hyperexcitability. **(A)** Näive DRG sensory neurons were incubated with either vehicle (Veh), 100 µM (CM) or 100 µM (SM) overnight before the effects on **(B)** resting membrane potential (RMP) and **(C)** rheobase were assessed by whole-cell patch clamp electrophysiology. *** *p-adj* < 0.001, **** *p-adj* < 0.0001: One-way ANOVA with Bonferroni-correct post hoc, n = 27 (Veh), 21 (CM), 26 (SM) cells from 8 mice (5 males (open symbols) and 3 females (closed symbols)).

## Supplementary Tables

**Table 1:**
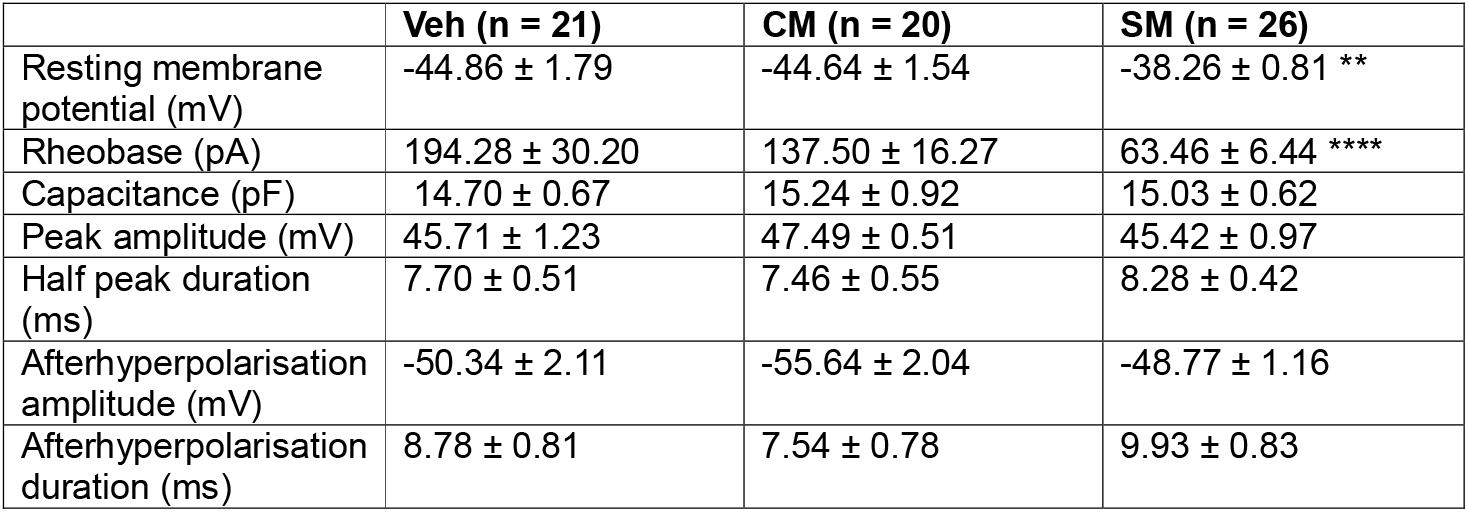
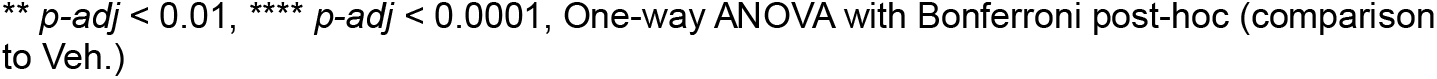
Intrinsic and active properties of naïve dorsal root ganglion neurons following overnight incubation with Veh, 1 µM CM or 1 µM SM.

|  | Veh (n = 21) | CM (n = 20) | SM (n = 26) |
| --- | --- | --- | --- |
| Resting membrane potential (mV) | -44.86 ± 1.79 | -44.64 ± 1.54 | -38.26 ± 0.81 ** |
| Rheobase (pA) | 194.28 ± 30.20 | 137.50 ± 16.27 | 63.46 ± 6.44 **** |
| Capacitance (pF) | 14.70 ± 0.67 | 15.24 ± 0.92 | 15.03 ± 0.62 |
| Peak amplitude (mV) | 45.71 ± 1.23 | 47.49 ± 0.51 | 45.42 ± 0.97 |
| Half peak duration (ms) | 7.70 ± 0.51 | 7.46 ± 0.55 | 8.28 ± 0.42 |
| Afterhyperpolarisation amplitude (mV) | -50.34 ± 2.11 | -55.64 ± 2.04 | -48.77 ± 1.16 |
| Afterhyperpolarisation duration (ms) | 8.78 ± 0.81 | 7.54 ± 0.78 | 9.93 ± 0.83 |

\*\* $p\text{-adj} < 0.01$ , \*\*\*\* $p\text{-adj} < 0.0001$ , One-way ANOVA with Bonferroni post-hoc (comparison to Veh.)

**Table 2:** Intrinsic and active properties of naïve dorsal root ganglion neurons following overnight incubation with Veh, 100 µM CM or 100 µM SM.

|  | Veh (n = 27) | CM (n = 21) | SM (n = 26) |
| --- | --- | --- | --- |
| Resting membrane potential (mV) | -61.1 ± 0.74 | -50.9 ± 1.51 **** | -43.4 ± 1.57 **** |
| Rheobase (pA) | 692.0 ± 49.8 | 395.0 ± 51.3 *** | 394.0 ± 45.6 **** |
| Capacitance (pF) | 27.69 ± 1.52 | 24.75 ± 1.76 | 25.38 ± 1.77 |
| Peak amplitude (mV) | 49.14 ± 0.92 | 50.28 ± 1.34 | 48.85 ± 1.11 |
| Half peak duration (ms) | 3.08 ± 0.26 | 4.94 ± 0.64 | 5.72 ± 0.75 ** |
| Afterhyperpolarisation amplitude (mV) | -65.31 ± 1.57 | -56.84 ± 1.14 *** | -50.10 ± 1.49 **** |
| Afterhyperpolarisation duration (ms) | 5.41 ± 0.50 | 6.60 ± 0.86 | 5.83 ± 0.47 |

\*\*\* $p\text{-adj} < 0.001$ , \*\*\*\* $p\text{-adj} < 0.0001$ , One-way ANOVA with Bonferroni post-hoc (comparison to Veh.)

**Table 3:** Intrinsic and active properties of knee-innervating dorsal root ganglion neurons from Veh-, CM- or SM-injected knee.

|  | Veh (n = 19) | CM (n = 37) | SM (n = 38) |
| --- | --- | --- | --- |
| Resting membrane potential (mV) | -51.7 ± 1.36 | -53.5 ± 0.82 | -47.3 ± 0.76 ** |
| Rheobase (pA) | 220.0 ± 31.4 | 236.0 ± 24.7 | 118.0 ± 14.2 * |
| Capacitance (pF) | 17.50 ± 1.18 | 15.76 ± 0.75 | 16.53 ± 0.81 |
| Peak amplitude (mV) | 55.51 ± 1.01 | 52.31 ± 0.95 | 53.49 ± 0.81 |
| Half peak duration (ms) | 6.18 ± 0.75 | 6.13 ± 0.59 | 7.53 ± 0.64 |
| Afterhyperpolarisation amplitude (mV) | -59.67 ± 1.29 | -60.74 ± 0.82 | -57.51 ± 1.08 |
| Afterhyperpolarisation duration (ms) | 7.52 ± 0.71 | 8.50 ± 0.75 | 8.64 ± 0.59 |

\* $p\text{-adj} < 0.05$ , \*\* $p\text{-adj} < 0.01$ , One-way ANOVA with Bonferroni post-hoc (comparison to Veh.)

